# Neutralization sensitivity of Omicron BA.2.75 to therapeutic monoclonal antibodies

**DOI:** 10.1101/2022.07.14.500041

**Authors:** Daichi Yamasoba, Izumi Kimura, Yusuke Kosugi, Keiya Uriu, Shigeru Fujita, Jumpei Ito, Kei Sato, The Genotype to Phenotype Japan (G2P-Japan) Consortium

## Abstract

Since the end of 2021, severe acute respiratory syndrome coronavirus 2 (SARS-CoV-2) Omicron variant outcompeted other variants and took over the world. After the emergence of original Omicron BA.1, Omicron BA.2 subvariant emerged and outcompeted BA.1. As of July 2022, some BA.2 subvariants, including BA.2.12.1, BA.4 and BA.5, emerged in multiple countries and begun outcompeting original BA.2. Moreover, a novel BA.2 subvariant, BA.2.75, was detected in eight countries including India at the end of June 2022, and preliminary investigations suggest that BA.2.75 is more transmissible over the other BA.2 subvariants. On July 7, 2022, the WHO classified BA.2.75 as a variant-of-concern lineage under monitoring. We have recently demonstrated that BA.4/5 is highly resistant to a therapeutic monoclonal antibody, cilgavimab, than BA.2. The resistance of SARS-CoV-2 variants to therapeutic antibodies can be attributed to the mutations in the viral spike protein. Compared to the BA.2 spike, BA.2.12.1 and BA.4/5 respectively bear two and four mutations in their spike proteins. On the other hand, the majority of BA.2.75 spike bears nine substitutions. The fact that the mutation number in the BA.2.75 spike is larger than those in the BA.4/5 spike raises the possibility that the BA.2.75 spike significantly reduces sensitivity towards therapeutic monoclonal antibodies than BA.2 and BA.4/5. In this study, we generated pseudoviruses harboring the spike proteins of BA.2.75, BA.4/5 and BA.2 and evaluated the efficacy of ten therapeutic monoclonal antibodies and three antibody cocktails against BA.2.75.

## Text

Since the end of 2021, severe acute respiratory syndrome coronavirus 2 (SARS-CoV-2) Omicron variant outcompeted other variants and took over the world. After the emergence of original Omicron BA.1, Omicron BA.2 subvariant emerged and outcompeted BA.1. As of July 2022, some BA.2 subvariants, including BA.2.12.1, BA.4 and BA.5, emerged in multiple countries and begun outcompeting original BA.2. Moreover, a novel BA.2 subvariant, BA.2.75, was detected in eight countries including India at the end of June 2022, and preliminary investigations suggest that BA.2.75 is more transmissible over the other BA.2 subvariants.^1^ On July 7, 2022, the WHO classified BA.2.75 as a variant-of-concern lineage under monitoring.^2^

We have recently demonstrated that BA.4/5 is highly resistant to a therapeutic monoclonal antibody, cilgavimab, than BA.2.^3^ The resistance of SARS-CoV-2 variants to therapeutic antibodies can be attributed to the mutations in the viral spike protein. Compared to the BA.2 spike, BA.2.12.1 and BA.4/5 respectively bear two and four mutations in their spike proteins.^3^ On the other hand, the majority of BA.2.75 spike bears nine substitutions (**Figure S1**). The fact that the mutation number in the BA.2.75 spike is larger than those in the BA.4/5 spike raises the possibility that the BA.2.75 spike significantly reduces sensitivity towards therapeutic monoclonal antibodies than BA.2 and BA.4/5. To address this possibility, we generated pseudoviruses harboring the spike proteins of BA.2.75, BA.4/5 and BA.2 and prepared ten therapeutic monoclonal antibodies and three antibody cocktails. Adintrevimab, bamlanivimab, casirivimab, etesevimab, and imdevimab did not work against BA.2, BA.4/5 and BA.2.75 (**Table 1** and **Figure S2**). Importantly, while regdanvimab, sotrovimab, and tixagevimab did not exhibit antiviral effects against BA.2 and BA.4/5, these three antibodies were functional against BA.2.75 (**Table 1**), suggesting that these antibodies can be used for the therapy and prevention of BA.2.75 infection. Consistent with our recent study,^3^ cilgavimab was less effective against BA.4/5 than BA.2, and BA.2.75 exhibited 24.4-fold higher resistance to cilgavimab than BA.2 [family-wise error rate (FWER)=0.04] (**Table 1**). Notably, although bebtelovimab exhibited robust antiviral effect against BA.2 and BA.4/5,^3^ BA.2.75 was significantly more resistant to this antibody than BA.2 (21.2-fold, FWER=0.01) and BA.4/5 (25.6-fold, FWER=0.01) (**Table 1**). These results suggest that bebtelovimab may not be a good choice to treat BA.2.75 infection.

**Table 1.** IC50s of ten therapeutic monoclonal antibodies against BA.2.75. Neutralization assay was performed using pseudoviruses harboring the SARS-CoV-2 spike proteins of BA.2, BA.4/5 (BA.2 spike:HV69-70del/L452R/F486V/R493Q) and BA.2.75 (BA.2 spike:K147E/W152R/F157L/I210V/G257S/D339H/G446S/N460K/R493Q) or the D614G-harboring B.1.1 lineage virus (B.1.1). Ten therapeutic monoclonal antibodies (adintrevimab, bamlanivimab, bebtelovimab, casirivimab, cilgavimab, etesevimab, imdevimab, regdanvimab, sotrovimab and tixagevimab) and three antibody cocktails [Ronapreve (casirivimab+imdevimab), Evusheld (cilgavimab+tixagevimab), and etesevimab+bamlanivimab] were tested. The assay of each antibody was performed in triplicate at each concentration to determine the 50% inhibitory concentration (IC50; ng/mL), and the assay was independently repeated four times. The presented data are expressed as the average ± 95% confidential interval. Statistical significance was evaluated by the Welch t-test with multiple testing corrections by the Holm method. An asterisk (*) and dagger (†) denote FWER < 0.05 for the BA.2.75 versus BA.2 and BA.2.75 versus BA.4/5 comparisons, respectively. Raw data and representative neutralization curves are respectively shown in **Table S1** and **Figure S2** in the Supplementary Appendix.

|  | B.1.1 | BA.2 | BA.4/5 | BA.2.75 |
| --- | --- | --- | --- | --- |
| Adintrevimab | 6.3 ± 1.7 | > 2750 | > 2750 | > 2750 |
| Bamlanivimab | 6.7 ± 1.1 | > 4725 | > 4725 | > 4725 |
| Bebtelovimab | 2.4 ± 0.9 | 1.7 ± 0.8 | 1.3 ± 0.3 | 34 ± 6.9 *† |
| Casirivimab | 3.4 ± 1.2 | > 5042 | > 5042 | 2303 ± 2570 |
| Cilgavimab | 14 ± 1.7 | 21 ± 7.9 | 305 ± 127 | 479 ± 154 * |
| Etesevimab | 12 ± 1.6 | > 4600 | > 4600 | > 4600 |
| Imdevimab | 8.0 ± 3.1 | > 5000 | > 5000 | > 5000 |
| Regdanvimab | 1.0 ± 0.4 | > 4025 | > 4025 | 42 ± 14 *† |
| Sotrovimab | 47 ± 50 | 1213 ± 224 | 1149 ± 159 | 240 ± 56 *† |
| Tixagevimab | 1.5 ± 0.6 | 3815 ± 1032 | > 4375 | 45 ± 8.2 *† |
| Ronapreve (casirivimab+imdevimab) | 3.9 ± 2.3 | > 5000 | > 5000 | > 5000 |
| Evusheld (cilgavimab+tixagevimab) | 4.7 ± 1.1 | 42 ± 17 | 586 ± 193 | 113 ± 31 * |
| Etesevimab+bamlanivimab | 8.3 ± 1.0 | > 4600 | > 4600 | > 4600 |

Mutations are accumulated in the S proteins of newly emerging SARS-CoV-2 variants. Therefore, the rapid evaluation of the efficiency of therapeutic monoclonal antibodies against novel SARS-CoV-2 variants should be important.

