## Supplementary Appendix for "Neutralization sensitivity of Omicron BA.2.75 to therapeutic monoclonal antibodies"

### Table of Contents

**Contents** Page


| | |
|---|---|
| **Materials and Methods** | 2-3 |
| Structural analysis | |
| Cell culture | |
| Plasmid construction | |
| Preparation of monoclonal antibodies | |
| Neutralization assay | |
| **Figure S1.** Amino acid substitutions in the BA.2.75 spike. | 4 |
| **Figure S2.** Representative neutralization curves. | 5-6 |
| **Table S1.** Summary of IC50 values and fold changes. | 7-12 |
| **Table S2.** Primers used for the construction of SARS-CoV-2 S expression plasmids. | 13 |
| **Consortia** | 14 |
| **Acknowledgments** | 15 |
| **Supplemental References** | 16-17 |

### Materials and Methods

#### Structural analysis

To indicate the positions of nine substitutions detected in the BA.2.75 spike (K147E/W152R/F157L/I210V/G257S/D339H/G446S/N460K/R493Q) in **Figure S1**, the cryo-electron microscopy structure of BA.2 spike[1] was used. The nine residues mutated in the BA.2.75 spike were labeled using the PyMOL molecular graphics system v2.5.0 (Schrödinger).

#### Cell culture

HEK293T cells (a human embryonic kidney cell line; ATCC CRL-3216) and HOS-ACE2/TMPRSS2 cells (kindly provided by Dr. Kenzo Tokunaga),[2,3] a derivative of HOS cells (a human osteosarcoma cell line; ATCC CRL-1543) stably expressing human ACE2 and TMPRSS2, were maintained in Dulbecco's modified Eagle's medium (DMEM) (high glucose) (Wako, Cat# 044-29765) containing 10% fetal bovine serum (FBS) (Sigma-Aldrich Cat# 172012-500ML), 100 units penicillin and 100 ug/mL streptomycin (PS) (Sigma-Aldrich, Cat# P4333-100ML).

#### Plasmid construction

To construct the plasmids expressing anti-SARS-CoV-2 monoclonal antibodies (adintrevimab, bamlanivimab, bebtelovimab, casirivimab, cilgavimab, etesevimab, imdevimab, regdanvimab, sotrovimab and tixagevimab), the sequences of the variable regions of these antibodies were obtained from KEGG Drug Database (https://www.genome.jp/kegg/drug/) and The Structural Antibody Database (http://opig.stats.ox.ac.uk/webapps/newsabdab/sabdab/) and were artificially synthesized by Fasmac. The obtained coding sequences of the variable regions of the heavy and light chains were cloned into the pCAGGS vector containing the sequences of the human immunoglobulin 1 and kappa constant region (kindly provided by Dr. Hisashi Arase). Plasmids expressing the SARS-CoV-2 spike proteins of the parental D614G (B.1.1), Omicron BA.2 and BA.4/5 were prepared in our previous studies.[3-6] Plasmids expressing the spike protein of Omicron variants BA.2.75 (GISAID ID: EPI_ISL_13471039) were generated by site-directed overlap extension PCR using pC-SARS2-S BA.2[6] as the template and the primers listed in **Table S2**. The resulting PCR fragment was subcloned into the KpnI-NotI site of the pCAGGS vector[7] using In-Fusion® HD Cloning Kit (Takara, Cat# Z9650N). Nucleotide sequences were determined by DNA sequencing services (Eurofins), and the sequence data were analyzed by Sequencher v5.1 software (Gene Codes Corporation).

#### Preparation of monoclonal antibodies

Ten monoclonal antibodies (adintrevimab, bamlanivimab, bebtelovimab, casirivimab, cilgavimab, etesevimab, imdevimab, regdanvimab, sotrovimab and tixagevimab) were prepared as previously described.[4,8,9] Briefly, the pCAGGS vectors containing the sequences encoding the immunoglobulin heavy and light chains were cotransfected into HEK293T cells at 1:1 ratio using PEI Max (Polysciences, Cat# 24765-1). The culture medium was refreshed with DMEM (low glucose) (Wako, Cat# 041-29775) containing 10% FBS without PS. At 72 h posttransfection, the culture medium was harvested, and the antibodies were purified using NAb protein A plus spin kit (Thermo Fisher Scientific, Cat# 89948) according to the manufacturer’s protocol.

**Neutralization assay**

Pseudoviruses were prepared as previously described.[4-6,9-15] Briefly, lentivirus (HIV-1)-based, luciferase-expressing reporter viruses were pseudotyped with the SARS-CoV-2 spikes. HEK293T cells ($3 \times 10^6$ cells) were cotransfected with 4 μg psPAX2-IN/HiBiT,[16] 4 μg pWPI-Luc2,[16] and 2 μg plasmids expressing parental S or its derivatives using PEI Max (Polysciences, Cat# 24765-1) according to the manufacturer's protocol. Two days post transfection, the culture supernatants were harvested and centrifuged. The pseudoviruses were stored at –80°C until use.

Neutralization assays were performed as previously described.[4-6,11,13,14] Briefly, the SARS-CoV-2 spike pseudoviruses (counting ~25,000 relative light units) were incubated with serially diluted monoclonal antibodies at 37°C for 1 h. Pseudoviruses without monoclonal antibody were included as controls. Then, a 40 μl mixture of pseudovirus and serum was added to HOS-ACE2/TMPRSS2 cells (10,000 cells/50 μl) in a 96-well white plate. Two days post infection, the infected cells were lysed with a Bright-Glo luciferase assay system (Promega, Cat# E2620), and the luminescent signal was measured using a GloMax explorer multimode microplate reader 3500 (Promega). The assay of each monoclonal antibody was performed in triplicate, and the 50% inhibitory concentration was calculated using Prism 9 (GraphPad Software).

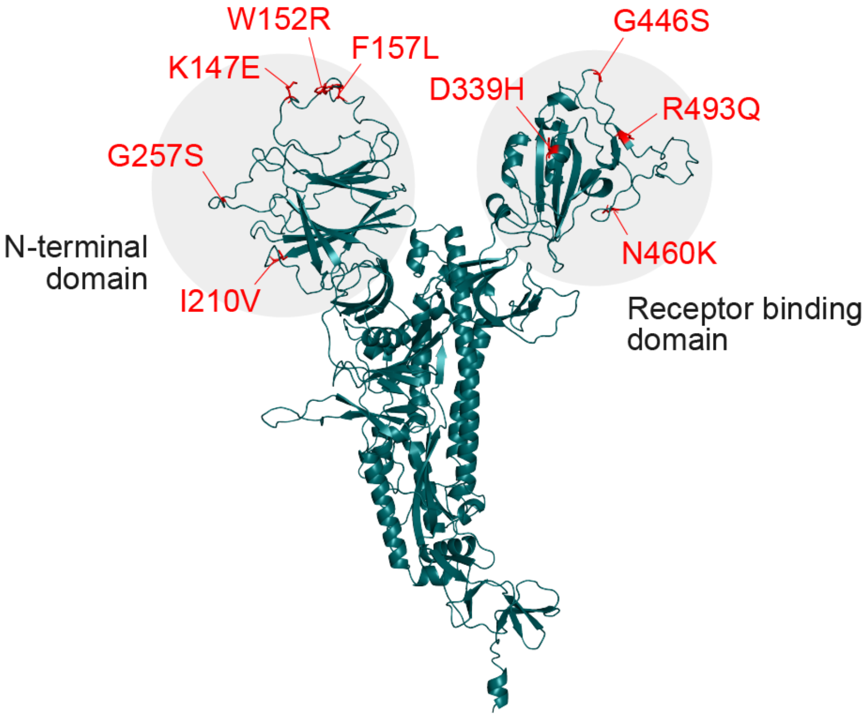


**Figure S1. Amino acid substitutions in the BA.2.75 spike.** In the cryo-electron microscopy structure of BA.2 spike,[1] the nine substitutions detected in the BA.2.75 spike are indicated in red. K147E, W152R, F157L, I210V, and G257S substitutions are located in the N-terminal domain (residues 13-304), while D339H, G446S, N460K, and R493Q substitutions are located in the receptor-binding domain (residues 319-541). In particular, G446S, N460K, and R493Q substitutions are located in the receptor-binding motif (residues 438-508).

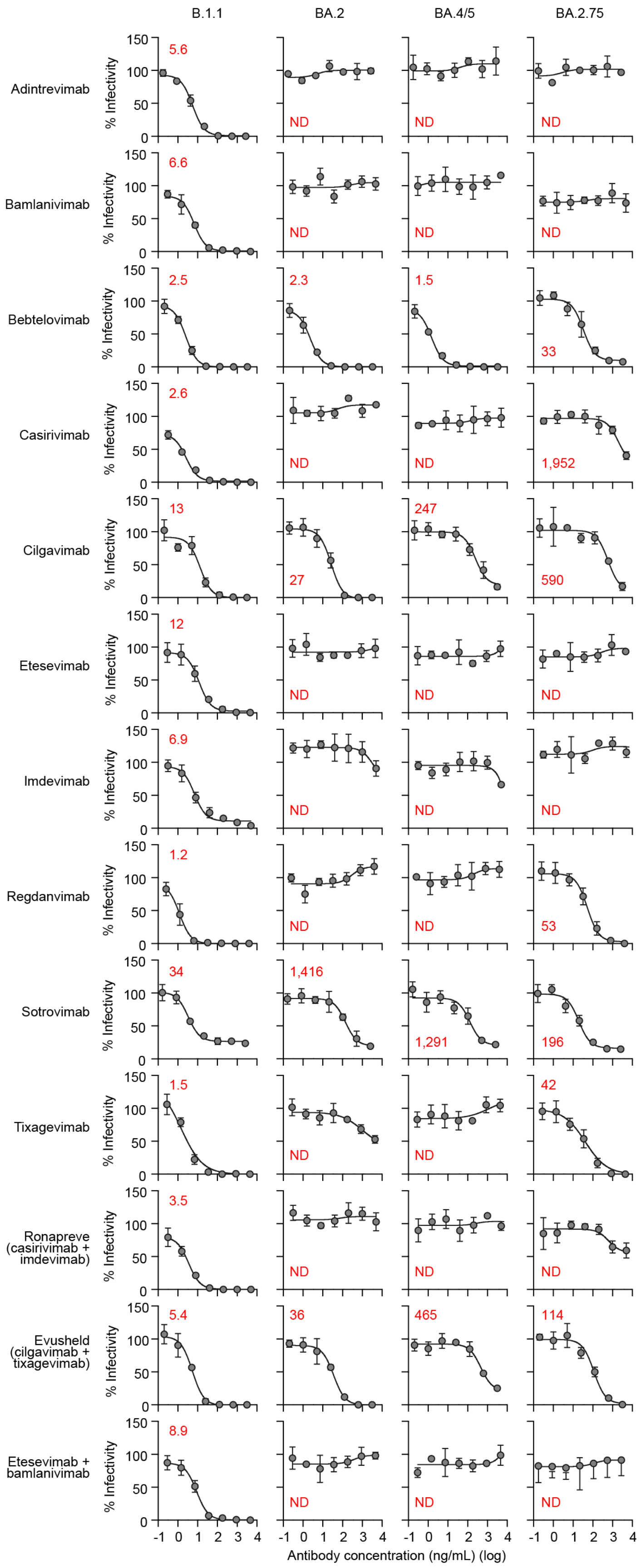
B.1.1
BA.2
BA.4/5
BA.2.75
Adintrevimab
5.6
ND
ND
ND
Bamlanivimab
6.6
ND
ND
ND
Bebtelovimab
2.5
2.3
1.5
33
Casirivimab
2.6
ND
ND
1,952
Cilgavimab
13
27
247
590
Etesevimab
12
ND
ND
ND
Imdevimab
6.9
ND
ND
ND
Regdanvimab
1.2
ND
ND
53
Sotrovimab
34
1,416
1,291
196
Tixagevimab
1.5
ND
ND
42
Ronapreve (casirivimab + imdevimab)
3.5
ND
ND
ND
Evusheld (cilgavimab + tixagevimab)
5.4
36
465
114
Etesevimab + bamlanivimab
8.9
ND
ND
ND
% Infectivity
0
50
100
150
-1 0 1 2 3 4
Antibody concentration (ng/mL) (log)

**Figure S2. Representative neutralization curves.** A neutralization assay was performed using pseudoviruses harboring the SARS-CoV-2 S proteins of BA.2 and BA.4/5 (BA.2 spike:HV69-70del/L452R/F486V/R493Q) and BA.2.75 (BA.2 spike:K147E/W152R/F157L/I210V/G257S/D339H/G446S/N460K/R493Q) or the D614G-harboring B.1.1 lineage virus (B.1.1). Ten therapeutic monoclonal antibodies (adintrevimab, bamlanivimab, bebtelovimab, casirivimab, cilgavimab, etesevimab, imdevimab, regdanvimab, sotrovimab and tixagevimab) and three antibody cocktails [Ronapreve (casirivimab+imdevimab), Evusheld (cilgavimab+tixagevimab), and etesevimab+bamlanivimab] were tested. The assay of each antibody was performed in triplicate at each concentration to determine the 50% inhibitory concentration (IC50; ng/mL), and the assay was independently repeated four times. Representative neutralization curves of a neutralization assay are shown. The red numbers in the panels indicate the IC50 value (ng/mL). ND, not determined. Summarized data of four independent assays are shown in **Table 1**. The IC50 value of each assay is summarized in **Table S1**.

**Table S1. Summary of IC50 values and fold changes.**

The value over the highest concentration is indicated in red.

**IC50 (ng/mL)**

| Adintrevimab | B.1.1 | BA.2 | BA.4/5 | BA.2.75 |
|---|---|---|---|---|
| Experiment 1 | 5.4 | >2750 | >2750 | >2750 |
| Experiment 2 | 7.8 | >2750 | >2750 | >2750 |
| Experiment 3 | 5.6 | >2750 | >2750 | >2750 |
| Experiment 4 | 6.3 | >2750 | >2750 | >2750 |

| Bamlanivimab | B.1.1 | BA.2 | BA.4/5 | BA.2.75 |
|---|---|---|---|---|
| Experiment 1 | 5.7 | >4725 | >4725 | >4725 |
| Experiment 2 | 7.2 | >4725 | >4725 | >4725 |
| Experiment 3 | 7.2 | >4725 | >4725 | >4725 |
| Experiment 4 | 6.6 | >4725 | >4725 | >4725 |

| Bebtelovimab | B.1.1 | BA.2 | BA.4/5 | BA.2.75 |
|---|---|---|---|---|
| Experiment 1 | 2.0 | 1.8 | 1.1 | 28 |
| Experiment 2 | 3.0 | 1.6 | 1.4 | 36 |
| Experiment 3 | 1.9 | 1.1 | 1.3 | 37 |
| Experiment 4 | 2.5 | 2.3 | 1.5 | 33 |

| Casirivimab | B.1.1 | BA.2 | BA.4/5 | BA.2.75 |
|---|---|---|---|---|
| Experiment 1 | 2.9 | >5042 | >5042 | 973 |
| Experiment 2 | 4.0 | >5042 | >5042 | 1639 |
| Experiment 3 | 2.6 | >5042 | >5042 | 1952 |
| Experiment 4 | 4.1 | >5042 | >5042 | 4647 |

| Cilgavimab | B.1.1 | BA.2 | BA.4/5 | BA.2.75 |
|---|---|---|---|---|
| Experiment 1 | 13 | 20 | 397 | 405 |
| Experiment 2 | 13 | 27 | 247 | 590 |
| Experiment 3 | 13 | 15 | 229 | 530 |
| Experiment 4 | 15 | 21 | 345 | 392 |

| Etesevimab | B.1.1 | BA.2 | BA.4/5 | BA.2.75 |
|---|---|---|---|---|
| Experiment 1 | 13 | >4600 | >4600 | >4600 |
| Experiment 2 | 12 | >4600 | >4600 | >4600 |
| Experiment 3 | 12 | >4600 | >4600 | >4600 |
| Experiment 4 | 11 | >4600 | >4600 | >4600 |

| Imdevimab | B.1.1 | BA.2 | BA.4/5 | BA.2.75 |
|---|---|---|---|---|
| Experiment 1 | 11 | >5000 | >5000 | >5000 |
| Experiment 2 | 7.9 | >5000 | >5000 | >5000 |
| Experiment 3 | 6.4 | >5000 | >5000 | >5000 |
| Experiment 4 | 6.9 | >5000 | >5000 | >5000 |

(Table S1, continued)

| Regdanvimab | B.1.1 | BA.2 | BA.4/5 | BA.2.75 |
|---|---|---|---|---|
| Experiment 1 | 0.8 | >4025 | >4025 | 32 |
| Experiment 2 | 1.2 | >4025 | >4025 | 53 |
| Experiment 3 | 1.0 | >4025 | >4025 | 38 |
| Experiment 4 | 1.3 | >4025 | >4025 | 43 |

| Sotrovimab | B.1.1 | BA.2 | BA.4/5 | BA.2.75 |
|---|---|---|---|---|
| Experiment 1 | 15.3 | 1141 | 1072 | 231 |
| Experiment 2 | 48.0 | 1194 | 1146 | 257 |
| Experiment 3 | 33.6 | 1416 | 1291 | 196 |
| Experiment 4 | 89.3 | 1100 | 1087 | 277 |

| Tixagevimab | B.1.1 | BA.2 | BA.4/5 | BA.2.75 |
|---|---|---|---|---|
| Experiment 1 | 1.1 | >4375 | >4375 | 48 |
| Experiment 2 | 1.5 | >4375 | >4375 | 42 |
| Experiment 3 | 1.6 | 3313 | >4375 | 32 |
| Experiment 4 | 2.0 | 3197 | >4375 | 59 |

| Ronapreve (casirivimab + imdevimab) | B.1.1 | BA.2 | BA.4/5 | BA.2.75 |
|---|---|---|---|---|
| Experiment 1 | 2.0 | >5000 | >5000 | >5000 |
| Experiment 2 | 4.6 | >5000 | >5000 | >5000 |
| Experiment 3 | 3.5 | >5000 | >5000 | >5000 |
| Experiment 4 | 5.4 | >5000 | >5000 | >5000 |

| Evusheld (cilgavimab + tixagevimab) | B.1.1 | BA.2 | BA.4/5 | BA.2.75 |
|---|---|---|---|---|
| Experiment 1 | 4.1 | 32 | 657 | 141 |
| Experiment 2 | 5.4 | 36 | 465 | 114 |
| Experiment 3 | 4.2 | 46 | 502 | 101 |
| Experiment 4 | 5.2 | 56 | 718 | 97 |

| Etesevimab + bamlanivimab | B.1.1 | BA.2 | BA.4/5 | BA.2.75 |
|---|---|---|---|---|
| Experiment 1 | 8.9 | >4600 | >4600 | >4600 |
| Experiment 2 | 8.4 | >4600 | >4600 | >4600 |
| Experiment 3 | 8.5 | >4600 | >4600 | >4600 |
| Experiment 4 | 7.4 | >4600 | >4600 | >4600 |

(Table S1, continued)

**Fold change (versus B.1.1)**

| Adintrevimab | B.1.1 | BA.2 | BA.4/5 | BA.2.75 |
|---|---|---|---|---|
| Experiment 1 | 1.0 | 508 | 508 | 508 |
| Experiment 2 | 1.0 | 354 | 354 | 354 |
| Experiment 3 | 1.0 | 494 | 494 | 494 |
| Experiment 4 | 1.0 | 433 | 433 | 433 |

| Bamlanivimab | B.1.1 | BA.2 | BA.4/5 | BA.2.75 |
|---|---|---|---|---|
| Experiment 1 | 1.0 | 823 | 823 | 823 |
| Experiment 2 | 1.0 | 655 | 655 | 655 |
| Experiment 3 | 1.0 | 660 | 660 | 660 |
| Experiment 4 | 1.0 | 711 | 711 | 711 |

| Bebtelovimab | B.1.1 | BA.2 | BA.4/5 | BA.2.75 |
|---|---|---|---|---|
| Experiment 1 | 1.0 | 0.9 | 0.6 | 14 |
| Experiment 2 | 1.0 | 0.5 | 0.5 | 12 |
| Experiment 3 | 1.0 | 0.6 | 0.7 | 20 |
| Experiment 4 | 1.0 | 0.9 | 0.6 | 13 |

| Casirivimab | B.1.1 | BA.2 | BA.4/5 | BA.2.75 |
|---|---|---|---|---|
| Experiment 1 | 1.0 | 1742 | 1742 | 336 |
| Experiment 2 | 1.0 | 1269 | 1269 | 413 |
| Experiment 3 | 1.0 | 1941 | 1941 | 751 |
| Experiment 4 | 1.0 | 1217 | 1217 | 1122 |

| Cilgavimab | B.1.1 | BA.2 | BA.4/5 | BA.2.75 |
|---|---|---|---|---|
| Experiment 1 | 1.0 | 1.5 | 29 | 30 |
| Experiment 2 | 1.0 | 2.0 | 18 | 44 |
| Experiment 3 | 1.0 | 1.2 | 18 | 42 |
| Experiment 4 | 1.0 | 1.4 | 23 | 26 |

| Etesevimab | B.1.1 | BA.2 | BA.4/5 | BA.2.75 |
|---|---|---|---|---|
| Experiment 1 | 1.0 | 351 | 351 | 351 |
| Experiment 2 | 1.0 | 386 | 386 | 386 |
| Experiment 3 | 1.0 | 387 | 387 | 387 |
| Experiment 4 | 1.0 | 436 | 436 | 436 |

| Imdevimab | B.1.1 | BA.2 | BA.4/5 | BA.2.75 |
|---|---|---|---|---|
| Experiment 1 | 1.0 | 466 | 466 | 466 |
| Experiment 2 | 1.0 | 630 | 630 | 630 |
| Experiment 3 | 1.0 | 784 | 784 | 784 |
| Experiment 4 | 1.0 | 721 | 721 | 721 |

(Table S1, continued)

| Regdanvimab | B.1.1 | BA.2 | BA.4/5 | BA.2.75 |
|---|---|---|---|---|
| Experiment 1 | 1.0 | 5357 | 5357 | 43 |
| Experiment 2 | 1.0 | 3326 | 3326 | 44 |
| Experiment 3 | 1.0 | 4226 | 4226 | 40 |
| Experiment 4 | 1.0 | 3137 | 3137 | 34 |

| Sotrovimab | B.1.1 | BA.2 | BA.4/5 | BA.2.75 |
|---|---|---|---|---|
| Experiment 1 | 1.0 | 74 | 70 | 15 |
| Experiment 2 | 1.0 | 25 | 24 | 5.4 |
| Experiment 3 | 1.0 | 42 | 38 | 5.8 |
| Experiment 4 | 1.0 | 12 | 12 | 3.1 |

| Tixagevimab | B.1.1 | BA.2 | BA.4/5 | BA.2.75 |
|---|---|---|---|---|
| Experiment 1 | 1.0 | 3941 | 3941 | 43 |
| Experiment 2 | 1.0 | 2956 | 2956 | 28 |
| Experiment 3 | 1.0 | 2137 | 2823 | 21 |
| Experiment 4 | 1.0 | 1567 | 2145 | 29 |

| Ronapreve (casirivimab + imdevimab) | B.1.1 | BA.2 | BA.4/5 | BA.2.75 |
|---|---|---|---|---|
| Experiment 1 | 1.0 | 2449 | 2449 | 2449 |
| Experiment 2 | 1.0 | 1083 | 1083 | 1083 |
| Experiment 3 | 1.0 | 1422 | 1422 | 1422 |
| Experiment 4 | 1.0 | 931 | 931 | 931 |

| Evusheld (cilgavimab + tixagevimab) | B.1.1 | BA.2 | BA.4/5 | BA.2.75 |
|---|---|---|---|---|
| Experiment 1 | 1.0 | 8.0 | 162 | 35 |
| Experiment 2 | 1.0 | 6.5 | 85 | 21 |
| Experiment 3 | 1.0 | 11 | 119 | 24 |
| Experiment 4 | 1.0 | 11 | 139 | 19 |

| Etesevimab + bamlanivimab | B.1.1 | BA.2 | BA.4/5 | BA.2.75 |
|---|---|---|---|---|
| Experiment 1 | 1.0 | 514 | 514 | 514 |
| Experiment 2 | 1.0 | 550 | 550 | 550 |
| Experiment 3 | 1.0 | 540 | 540 | 540 |
| Experiment 4 | 1.0 | 619 | 619 | 619 |

(Table S1, continued)

**Fold change (versus BA.2)**

| Adintrevimab | B.1.1 | BA.2 | BA.4/5 | BA.2.75 |
|---|---|---|---|---|
| Experiment 1 | 0.0020 | 1.0 | 1.0 | 1.0 |
| Experiment 2 | 0.0028 | 1.0 | 1.0 | 1.0 |
| Experiment 3 | 0.0020 | 1.0 | 1.0 | 1.0 |
| Experiment 4 | 0.0023 | 1.0 | 1.0 | 1.0 |

| Bamlanivimab | B.1.1 | BA.2 | BA.4/5 | BA.2.75 |
|---|---|---|---|---|
| Experiment 1 | 0.0012 | 1.0 | 1.0 | 1.0 |
| Experiment 2 | 0.0015 | 1.0 | 1.0 | 1.0 |
| Experiment 3 | 0.0015 | 1.0 | 1.0 | 1.0 |
| Experiment 4 | 0.0014 | 1.0 | 1.0 | 1.0 |

| Bebtelovimab | B.1.1 | BA.2 | BA.4/5 | BA.2.75 |
|---|---|---|---|---|
| Experiment 1 | 1.1 | 1.0 | 0.6 | 15 |
| Experiment 2 | 1.9 | 1.0 | 0.9 | 22 |
| Experiment 3 | 1.7 | 1.0 | 1.1 | 33 |
| Experiment 4 | 1.1 | 1.0 | 0.6 | 14 |

| Casirivimab | B.1.1 | BA.2 | BA.4/5 | BA.2.75 |
|---|---|---|---|---|
| Experiment 1 | 0.0006 | 1.0 | 1.0 | 0.2 |
| Experiment 2 | 0.0008 | 1.0 | 1.0 | 0.3 |
| Experiment 3 | 0.0005 | 1.0 | 1.0 | 0.4 |
| Experiment 4 | 0.0008 | 1.0 | 1.0 | 0.9 |

| Cilgavimab | B.1.1 | BA.2 | BA.4/5 | BA.2.75 |
|---|---|---|---|---|
| Experiment 1 | 0.69 | 1.0 | 20 | 21 |
| Experiment 2 | 0.50 | 1.0 | 9.2 | 22 |
| Experiment 3 | 0.86 | 1.0 | 16 | 36 |
| Experiment 4 | 0.73 | 1.0 | 17 | 19 |

| Etesevimab | B.1.1 | BA.2 | BA.4/5 | BA.2.75 |
|---|---|---|---|---|
| Experiment 1 | 0.0029 | 1.0 | 1.0 | 1.0 |
| Experiment 2 | 0.0026 | 1.0 | 1.0 | 1.0 |
| Experiment 3 | 0.0026 | 1.0 | 1.0 | 1.0 |
| Experiment 4 | 0.0023 | 1.0 | 1.0 | 1.0 |

| Imdevimab | B.1.1 | BA.2 | BA.4/5 | BA.2.75 |
|---|---|---|---|---|
| Experiment 1 | 0.0021 | 1.0 | 1.0 | 1.0 |
| Experiment 2 | 0.0016 | 1.0 | 1.0 | 1.0 |
| Experiment 3 | 0.0013 | 1.0 | 1.0 | 1.0 |
| Experiment 4 | 0.0014 | 1.0 | 1.0 | 1.0 |

(Table S1, continued)

| Regdanvimab | B.1.1 | BA.2 | BA.4/5 | BA.2.75 |
|---|---|---|---|---|
| Experiment 1 | 0.0002 | 1.0 | 1.0 | 0.008 |
| Experiment 2 | 0.0003 | 1.0 | 1.0 | 0.013 |
| Experiment 3 | 0.0002 | 1.0 | 1.0 | 0.009 |
| Experiment 4 | 0.0003 | 1.0 | 1.0 | 0.011 |

| Sotrovimab | B.1.1 | BA.2 | BA.4/5 | BA.2.75 |
|---|---|---|---|---|
| Experiment 1 | 0.013 | 1.0 | 0.9 | 0.2 |
| Experiment 2 | 0.040 | 1.0 | 1.0 | 0.2 |
| Experiment 3 | 0.024 | 1.0 | 0.9 | 0.1 |
| Experiment 4 | 0.081 | 1.0 | 1.0 | 0.3 |

| Tixagevimab | B.1.1 | BA.2 | BA.4/5 | BA.2.75 |
|---|---|---|---|---|
| Experiment 1 | 0.0003 | 1.0 | 1.0 | 0.011 |
| Experiment 2 | 0.0003 | 1.0 | 1.0 | 0.009 |
| Experiment 3 | 0.0005 | 1.0 | 1.3 | 0.010 |
| Experiment 4 | 0.0006 | 1.0 | 1.4 | 0.018 |

| Ronapreve (casirivimab + imdevimab) | B.1.1 | BA.2 | BA.4/5 | BA.2.75 |
|---|---|---|---|---|
| Experiment 1 | 0.0004 | 1.0 | 1.0 | 1.0 |
| Experiment 2 | 0.0009 | 1.0 | 1.0 | 1.0 |
| Experiment 3 | 0.0007 | 1.0 | 1.0 | 1.0 |
| Experiment 4 | 0.0011 | 1.0 | 1.0 | 1.0 |

| Evusheld (cilgavimab + tixagevimab) | B.1.1 | BA.2 | BA.4/5 | BA.2.75 |
|---|---|---|---|---|
| Experiment 1 | 0.1 | 1.0 | 20.2 | 4.3 |
| Experiment 2 | 0.2 | 1.0 | 13.1 | 3.2 |
| Experiment 3 | 0.1 | 1.0 | 11.0 | 2.2 |
| Experiment 4 | 0.1 | 1.0 | 12.8 | 1.7 |

| Etesevimab + bamlanivimab | B.1.1 | BA.2 | BA.4/5 | BA.2.75 |
|---|---|---|---|---|
| Experiment 1 | 0.0019 | 1.0 | 1.0 | 1.0 |
| Experiment 2 | 0.0018 | 1.0 | 1.0 | 1.0 |
| Experiment 3 | 0.0019 | 1.0 | 1.0 | 1.0 |
| Experiment 4 | 0.0016 | 1.0 | 1.0 | 1.0 |

**Table S2. Primers used for the construction of SARS-CoV-2 S expression plasmids.**

| Primer name | Sequence (5'-to-3') |
|---|---|
| Omicron universal Fw | cactatagggcgaattgggtaccatgtttgtgttcctggt |
| BA2 Rv | agctccaccgcggtggcggccgctcaggtgtagtgcagtttca |
| pC_S_BA2_147_152_157_R | cagactccattctggacttgttgttctcgtggtagtagac |
| pC_S_BA2_147_152_157_F | caacaagtccagaatggagtctgagctgagggtctactcc |
| pC_S_BA2_I210V_R | cctgcccaggttcactggtgtgtgttt |
| pC_S_BA2_I210V_F | aaacacacaccagtgaacctgggcagg |
| pC_S_BA2_G257S_R | tcctgctgtccagctagaggaggagtc |
| pC_S_BA2_G257S_F | gactcctcctctagctggacagcagga |
| pC_S_BA2_D339H_R | attgaacacctcgtgaaatggacacag |
| pC_S_BA2_D339H_F | ctgtgtccatttcacgaggtgttcaat |
| pC_S_BA2_G446S_R | gttgtagttgccgctcaccttgctgtc |
| pC_S_BA2_G446S_F | gacagcaaggtgagcggcaactacaac |
| pC_S_BA2_N460K_R | aaatggtttcagcttgctcttcctgaa |
| pC_S_BA2_N460K_F | ttcaggaagagcaagctgaaaccattt |
| pC-S_BA2_R493Q-F | ttactttccactccaatcctatggcttca |
| pC-S_BA2_R493Q-R | tgaagccataggattggagtggaaagtaa |

### Consortia

#### The Genotype to Phenotype Japan (G2P-Japan) Consortium

**The Institute of Medical Science, The University of Tokyo, Japan**
Mai Suganami, Mika Chiba, Ryo Yoshimura, Naoko Misawa

**Hokkaido University, Japan**
Keita Matsuno, Naganori Nao, Hirofumi Sawa, Mai Kishimoto, Shinya Tanaka, Masumi Tsuda, Lei Wang, Yoshikata Oda, Marie Kato, Zannatul Ferdous, Hiromi Mouri, Kenji Shishido, Takasuke Fukuhara, Tomokazu Tamura, Rigel Suzuki, Hayato Ito

**Tokyo Metropolitan Institute of Public Health, Japan**
Kenji Sadamasu, Kazuhisa Yoshimura, Hiroyuki Asakura, Isao Yoshida, Mami Nagashima

**Tokai University, Japan**
So Nakagawa, Jiaqi Wu

**Kyoto University, Japan**
Akifumi Takaori-Kondo, Kotaro Shirakawa, Kayoko Nagata, Yasuhiro Kazuma, Ryosuke Nomura, Yoshihito Horisawa, Yusuke Tashiro, Yugo Kawai

**Hiroshima University, Japan**
Takashi Irie, Ryoko Kawabata

**Kumamoto University, Japan**
Terumasa Ikeda, Hesham Nasser, Ryo Shimizu, MST Monira Begum, Otowa Takahashi, Kimiko Ichihara, Takamasa Ueno, Chihiro Motozono, Mako Toyoda

**University of Miyazaki, Japan**
Akatsuki Saito, Erika P Butlertanaka, Yuri L Tanaka, Maya Shofa

**Acknowledgments**

We would like to thank all members of The Genotype to Phenotype Japan (G2P-Japan) Consortium. We thank Dr. Kenzo Tokunaga (National Institute of Infectious Diseases, Japan) and Dr. Hisashi Arase (Osaka University, Japan) for sharing materials. We gratefully acknowledge the numerous laboratories worldwide that have provided sequence data and metadata to GISAID. A full list of originating and submitting laboratories for the sequences used in our analysis can be found at https://www.gisaid.org using the EPI-SET-ID: EPI_SET_20220712sf.
